# De novo genome assemblies of the dwarf honey bee subgenus *Micrapis*: *Apis andreniformis* and *Apis florea*

**DOI:** 10.1101/2025.04.01.646657

**Authors:** Atma Ivancevic, Madison Sankovitz, Holly Allen, Olivia Joyner, Edward B. Chuong, Samuel D. Ramsey

**Author notes:** **Corresponding Authors:** Samuel D. Ramsey, 334 UCB, Boulder, CO 80309, Edward B. Chuong, 596 UCB, Boulder, CO 80309. co-first authors (Atma Ivancevic, Madison Sankovitz). co-last authors (Edward B. Chuong, Samuel D. Ramsey).

## Abstract

The *Micrapis* subgenus, which includes the black dwarf honey bee (*Apis andreniformis*) and the red dwarf honey bee (*Apis florea*), remains underrepresented in genomic studies despite its ecological significance. Here, we present high-quality de novo genome assemblies for both species, generated using a hybrid sequencing approach combining Oxford Nanopore Technologies (ONT) long reads with Illumina short reads. The final assemblies are highly contiguous, with contig N50 values of 5.0 Mb (*A. andreniformis*) and 4.3 Mb (*A. florea*), representing a major improvement over the previously published *A. florea* genome. Genome completeness assessments indicate high quality, with BUSCO scores exceeding 98.5% and k-mer analyses supporting base-level accuracy. Repeat annotation revealed a relatively low repetitive sequence content (∼6%), consistent with other *Apis* species. Using RNA sequencing data, we annotated 12,232 genes for *A. andreniformis* and 12,597 genes for *A. florea*, with >97% completeness in predicted proteomes. These genome assemblies provide a valuable resource for comparative and functional genomic studies, offering new insights into the genetic basis of dwarf honey bee adaptations.

## Introduction

Honey bees (*Apis* spp.) are among the most ecologically and economically significant insects on the planet (Hristov et al. 2020; Khalifa et al. 2021; Papa et al. 2022). Renowned for their role in pollination, honey production, and as models for studying social behavior, these eusocial insects are indispensable to agricultural systems and natural ecosystems (Gill 1990; Zayed and Robinson 2012; Hoover and Ovinge 2018). There are eight recognized honey bee species that exhibit varying sizes, nest structures, and ecological niches, though the total number of species is disputed, with some evidence for 11 (Crane 2009). While the western honey bee *A. mellifera* is the most extensively studied and globally distributed species, other honey bee species, including those in the subgenera *Apis, Megapis*, and *Micrapis*, exhibit remarkable diversity in their behaviors and adaptations, many of which remain underexplored.

The subgenus *Micrapis* consists of the dwarf honey bees (**Fig. 1**), black *A. andreniformis* and red *A. florea*, which live throughout south and southeast Asia (Benjamin P. Oldroyd and Wongsiri 2009). Red dwarf honey bees have been shown to contribute to the pollination of a diverse range of flora, playing an essential role in maintaining the biodiversity and ecosystem stability of its native regions (Ali, Sajjad, and Saeed 2017; Abrol 2010; Shwetha, Bhat, and Neethu 2020). Black dwarf honey bees are less studied, likely because they have a smaller distribution than their sister species, although their size and life history are similar and therefore they likely have a comparable ecological impact (Otis 1996). However, despite their ecological importance, both dwarf honey bee species remain underexplored compared to the widely studied western honey bee.

**Fig. 1.**
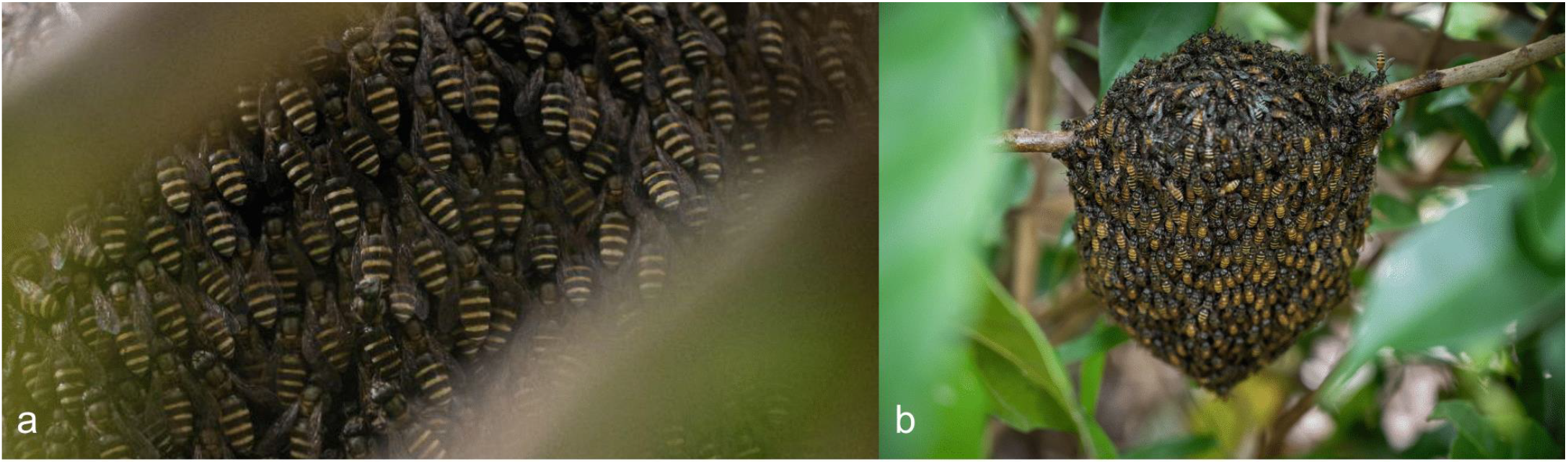
The dwarf honey bees: (a) black dwarf honey bees and (b) red dwarf honey bees. Photos by Sirachai Shin Arunrugstichai.

Dwarf honey bees are morphologically and behaviorally distinct from western honey bees. Dwarf honey bees only live in specific regions in Asia, whereas western honey bees have a cosmopolitan distribution (Otis 1996). Workers of the dwarf honey bees are significantly smaller, about half the size of western honey bee workers (B. P. Oldroyd 2021). They also exhibit smaller colony sizes, a simplicity that extends to their nesting behavior; both species construct single, exposed combs in open-air settings, typically hanging from thin branches of low shrubs (Rinderer et al. 1996; B. P. Oldroyd 2021). This contrasts with the enclosed nests of western honey bees (Seeley and Morse 1976).

Comparative genomics across honey bee species is critical for understanding the genetic basis of their distinctive traits, including morphology and ecological adaptations. Individual genomes of the western honey bee *Apis mellifera* have led to the identification of new subspecies, a better understanding of evolutionary lineages, and the discovery of a significant number of genes involved with adaptations and colony-level quantitative traits (Dogantzis and Zayed 2019). Genomics can dramatically improve our ability to understand invasive honey bee parasites and mitigate their impacts by selecting stocks with resilient traits, revealing the effect of parasites on bee health, and identifying biomarkers for rapid diagnostics (Grozinger and Zayed 2020). The dwarf honey bees have relatively few parasites that are otherwise invasive and highly damaging to western honey bee colonies worldwide; they are not hosts of *Varroa* mites, and *Tropilaelaps* mites have only been observed in red dwarf honey bee colonies in India several decades ago (Abrol and Kakroo 1997). Therefore, dwarf honey bee genomic resources are timely and may be valuable in our collective efforts to control invasive parasites. While a high-quality nuclear genome assembly is available for the western honey bee (Wallberg et al. 2019), the genomes of the dwarf honey bees remain poorly characterized. The nuclear genome of the red dwarf honey bee *A. florea* was previously sequenced using legacy 454 sequencing technology (Fouks et al. 2021), this assembly has relatively low coverage (20.5x) and is highly fragmented (total length: 211.3 Mb, scaffold N50: 2.9 Mb), making it suboptimal for genomic analysis. The nuclear genome of the black dwarf honey bee *A. andreniformis* has not been sequenced. To address this gap, we combined Oxford Nanopore Technologies (ONT) long-read sequencing with Illumina short-read sequencing to sequence both species’ nuclear genomes and transcriptomes, generating high-quality genome assemblies and transcript-level gene annotations for both dwarf honey bees. These assemblies provide a valuable resource for comparative genomic analyses and offer new insights into the genetic underpinnings of traits unique to dwarf honey bees.

## Materials and Methods

### Sample collection

Genome sequencing was performed using specimens of *A. andreniformis* and *A. florea* collected from managed colonies in Singapore. Specifically, one female worker *A. andreniformis* pupa (Aa1SG1, diploid) was collected from a colony in Frankel, and one female worker *A. florea* pupa (Af1SG1, diploid) was collected from a colony in Admiralty.

To complement genome sequencing, additional samples from the same colonies were collected for RNA sequencing. These included:

- Two female worker *A. andreniformis* pupae (diploid, samples Aa1SG2 and Aa1SG3).
- One female worker *A. florea* pupa (diploid, Af1SG2) and one male *A. florea* pupa (haploid, Af1SG3).

Pupae were extracted from comb cells with forceps after removing the wax cell capping with forceps. After collection, all specimens were flash-frozen in liquid nitrogen and stored at -80°C until extraction. They were then shipped in a cryo-shipper from Singapore to Colorado.

### DNA extraction and sequencing

Genomic DNA was extracted from one female *A. andreniformis* pupa and one female *A. florea* pupa using the Quick-DNA Tissue/Insect Kit (Zymo Research), according to the manufacturer’s instructions. For ONT long-read sequencing, a genomic library was prepared with the Native Barcoding Kit 24 V14 kit (SQK-NBD114.24) and sequenced on an R10.4.1 PromethION flow cell (FLO-PRO114M). For Illumina sequencing, a genomic library was prepared using the Ovation Ultralow System V2 kit (Tecan) and sequenced on an Illumina NovaSeq 6000 (University of Colorado Genomics Core) at the University of Colorado, Anschutz. Paired-end 2 × 150 bp reads were generated.

### RNA extraction and sequencing

RNA was extracted from two female *A. andreniformis* pupae, one female *A. florea* pupa, and one male *A. florea* pupa using a TRIzol/column-based method. In brief, whole pupa (30-110 mg) were homogenized in 1 ml TRIzol (ThermoFisher) and then centrifuged at 12,000 xg for 5 minutes at 4°C to remove the cell debris. The supernatant was incubated for 5 minutes at room temperature, mixed with 200 µL chloroform, incubated for a further 3 minutes, and then centrifuged at 12,000 xg for 5 minutes at 4°C. The upper, aqueous phase was combined with a 1:1 ratio of 100% ethanol and then purified according to the Direct-zol RNA Miniprep Kit (Zymo Research).

For ONT sequencing, an RNA library was prepared with the PCR cDNA Barcoding Kit (SQK-PCB111.24) and sequenced on an R9.4.1 PromethION flow cell (FLO-PRO002). For Illumina sequencing, an RNA library was prepared using the KAPA mRNA HyperPrep kit (KK8581) and sequenced on an Illumina NovaSeq 6000 (University of Colorado Genomics Core) at the University of Colorado, Anschutz. Paired-end 2 × 150 bp reads were generated.

### Genome assembly and polishing

High-quality genome assemblies for *A. andreniformis* and *A. florea* were generated using a combination of ONT long reads and Illumina short reads. The assembly process was conducted in several steps, including base-calling, demultiplexing, *de novo* assembly, initial polishing, and additional polishing using the short reads.

ONT raw signals in POD5 format were base-called using Dorado v0.7.2 (ONT Public License v1.0) using the super high accuracy model (SUP) to ensure high-quality basecalls, and --kit-name SQK-NBD114-24 to enable barcode classification in-line with basecalling. Reads were demultiplexed using Dorado’s demux function and flags --no-classify and --emit-fastq to produce FASTQ files for each barcode. Read statistics for demultiplexed reads were obtained using NanoStat v1.6.0 (De Coster et al. 2018).

De novo assemblies of the draft genomes for *A. andreniformis* and *A. florea* were conducted using Flye v2.9.5 (Kolmogorov et al. 2019) with options --nano-hq to indicate high-quality reads and the genome size estimate set to 230 Mb (--genome-size 230m) for both bees. The mean coverage of each genome was derived from Flye v2.9.5 (Kolmogorov et al. 2019). Draft assemblies were polished using Medaka v2.0.0 (ONT Public License v1.0), providing the raw ONT reads for each bee and the model r1041_e82_400bps_sup_v5.0.0 as input. Further polishing was conducted with Pilon v1.24 (Walker et al. 2014), leveraging Illumina short reads to correct small indels and base call mismatches.

Illumina short reads were first trimmed for adapter sequences, low-quality bases, and contaminants using the bbduk.sh script from BBMap v38.05 (https://sourceforge.net/projects/bbmap) and the following parameters: -Xmx32g ktrim=r k=31 mink=11 hdist=1 tpe tbo qtrim=r trimq=10. Trimmed reads were aligned to their corresponding long-read draft assemblies using BWA-MEM v0.7.5 (Li and Durbin 2009), with alignments filtered to exclude low-quality and unmapped reads (-q 10 -F 4) and sorted with Samtools v1.16.1 (Danecek et al. 2021; Li et al. 2009). The resulting BAM files were used in Pilon v1.24, as described above, to polish assemblies at the per-base level, correcting small base pair errors and indels. All software was executed using default settings unless otherwise specified.

After polishing, VSEARCH v2.14.1 (Rognes et al. 2016) was used with function --sortbylength to sort all contigs by length, and parameters --maxseqlength 1000000000 --minseqlength 3000 to remove contigs shorter than 3 kb for each genome. Before submission to GenBank, the final assemblies underwent contamination screening using NCBI’s Foreign Contamination Screen Tool Suite v0.5.4 (Astashyn et al. 2024). This process consisted of two sequential steps: (1) screening for vector and adapter contamination using FCS-adaptor, and (2) identifying and removing foreign contaminants such as bacterial and viral sequences using FCS-GX, with Apis-specific taxonomic filtering (NCBI taxonomy ID 7464 for *A. andreniformis* and taxonomy ID 7463 for *A. florea*). The cleaned assemblies were then submitted to GenBank.

### Quality assessment

The quality of each genome was assessed based on three criteria: assembly contiguity, genome completeness, and k-mer comparison to short reads.

Assembly contiguity was evaluated using QUAST v5.2.0 (Mikheenko et al. 2018) to generate metrics such as N50, L50, total assembly length, and the number of contigs.

Genome completeness was assessed using BUSCO v5.7.1 (Manni et al. 2021) with OrthoDB v10 (Kriventseva et al. 2019) to evaluate the presence or absence of universal single-copy orthologous genes. Detected BUSCO genes were classified as single-copy, duplicated, or fragmented. The following lineage-specific OrthoDB databases were used:

- **hymenoptera_odb10** (creation date: 2024-01-08; 40 genomes; 5991 BUSCOs)
- **insecta_odb10** (creation date: 2024-01-08; 75 genomes; 1367 BUSCOs)
- **arthropoda_odb10** (creation date: 2024-01-08; 90 genomes; 1013 BUSCOs)
- **metazoa_odb10** (creation date: 2024-01-08; 65 genomes; 954 BUSCOs)

Completeness statistics were compared to two existing *Apis* genome assemblies: *A. mellifera* Amel_HAv3.1 (NCBI RefSeq assembly accession GCF_003254395.2) (Wallberg et al. 2019) and *A. florea* Aflo_1.1 (NCBI RefSeq assembly accession GCF_000184785.3) (Fouks et al. 2021). K-mer analysis was performed using Meryl v1.4.1 (Rhie et al. 2020) and Merqury v1.3 (Rhie et al. 2020) to compare k-mers between the short-read data and polished genome assemblies. Meryl was used to generate k-mer databases from the Illumina paired-end short reads. Merqury was used to compare these k-mer databases to the genome assemblies, providing estimates of k-mer completeness (i.e. the percentage of short-read k-mers present in the assembly) and assembly quality values. While the short reads in this study were only used for genome polishing, this approach nonetheless highlighted the concordance between the Illumina data and the final assemblies. All software was executed using default settings.

### Repeat annotation

RepeatMasker v4.1.7 (Smit et al. 2013, www.repeatmasker.org) was used to softmask repeats in the polished assemblies and generate repeat statistics for each genome. RepeatMasker was configured with Tandem Repeats Finder v4.0.9 (Benson 1999), RMBlast v2.14.0 (www.repeatmasker.org/rmblast), and all nine library partitions from Dfam Database v3.8 (Storer et al. 2021). RepeatMasker was executed with default parameters except -pa 8 -species Eukaryota - noisy -xsmall.

### Transcriptome assembly and annotation

The transcriptomes of *A. andreniformis* and *A. florea* were characterized using long-read ONT cDNA sequencing data and short-read Illumina RNA sequencing data.

Long-read cDNA sequencing data was base-called using Dorado v0.7.2 (ONT Public License v1.0) with the super high accuracy (SUP) model and the --no-trim option to retain untrimmed reads necessary for cDNA-specific downstream analyses. Demultiplexing was performed using Dorado’s demux function with --no-trim enabled. Read quality and length distributions were assessed using NanoStat v1.6.0 (De Coster et al. 2018).

To refine the ONT reads, Pychopper v2.5.0 (ONT Public License v2.0) was used to identify and trim full-length cDNA reads, with -k PCB111 specified for the barcode kit. High-confidence full-length reads were merged with rescued reads to generate the final processed cDNA dataset. ONT reads from the male *A. florea* sample (Af1SG3) were excluded from further analyses due to poor sequencing quality, an N50 of only 196 bp, and high barcode retention. The remaining three samples—two *A. andreniformis* females and one *A. florea* female—demonstrated sufficient quality and coverage for downstream analyses, with N50 values exceeding 1,100 bp and total yields ranging from 31 to 40 Gb. Processed cDNA reads were aligned to the genome using Minimap2 v2.22 with the -x splice option.

Illumina RNA-seq reads were preprocessed using the bbduk.sh script from BBMap v38.05 (https://sourceforge.net/projects/bbmap) and the following parameters: -Xmx32g ktrim=r k=31 mink=11 hdist=1 tpe tbo qtrim=r trimq=10, to remove adapter sequences and low-quality bases. A Hisat2 genome index was generated for each species using Hisat2 v2.1.0 (Kim et al. 2019), and preprocessed reads were mapped to the genome assembly with default sensitivity and the --dta flag to optimize alignments for transcript assembly. BAM files were sorted and indexed using Samtools v1.16.1 (Danecek et al. 2021; Li et al. 2009), and the two Illumina RNA-seq BAM files were merged for each species.

Genome annotation was performed using BRAKER3 (Gabriel et al. 2024), an automated gene prediction pipeline that integrates RNA-seq and protein homology evidence to generate gene models. Gene prediction was carried out using GeneMark-ETP (Brůna, Lomsadze, and Borodovsky 2024) and AUGUSTUS (Stanke et al. 2008), incorporating extrinsic evidence to refine gene structures. TSEBRA (Gabriel et al. 2021) was used to select the most well-supported transcript isoforms. StringTie2 (Kovaka et al. 2019) was used for transcriptome assembly. To improve homology-based predictions, we provided the Arthropoda protein dataset from OrthoDB v12 (Tegenfeldt et al. 2025), downloaded from https://bioinf.uni-greifswald.de/bioinf/partitioned_odb12/Arthropoda.fa.gz. DIAMOND (Buchfink, Xie, and Huson 2015) was used to incorporate and align the Arthropoda protein sequences. Gene annotation outputs were further processed with GffRead (Pertea and Pertea 2020) for format conversion and filtering.

For *A. andreniformis*, BRAKER3 was run using two ONT cDNA BAM files (one per sample), a single merged Illumina RNA-seq BAM file, and protein sequences from Arthropoda as extrinsic evidence. For *A. florea*, BRAKER3 was run with one ONT cDNA BAM file (from sample Af1SG2), a single merged Illumina RNA-seq BAM file, and protein sequences. All software was executed using default settings unless otherwise specified.

## Results and Discussion

### Genome sequence data summary

Sequencing statistics are summarized in **Table 1**. In brief, for *A. andreniformis*, long-read sequencing using ONT (Native Barcoding V14 Kit sequenced on R10.4.1 PromethION flow cells) yielded 54.0 Gb of raw reads (read length N50 of 6,657 bp; mean read quality of 15.9). Using an estimated genome size of ∼230 Mb gave an initial coverage estimate of 235x. De novo assembly using Flye (Kolmogorov et al. 2019) resulted in a reported mean coverage of 235x. Whole genome sequencing using Illumina yielded 16.4 Gb (54,425,614 150 bp read pairs; mean quality score of 38.5), producing an estimated short read coverage of 71x. Pilon (Walker et al. 2014) was used to polish the long-read genome assembly with these Illumina short reads. All raw sequencing reads were uploaded to the NCBI Sequence Read Archive (SRA), which revealed that the ONT reads contained <0.01% bacteria and other contaminants. In contrast, the Illumina reads contained many unidentified reads (32.43%, likely derived from adapter sequences) and contaminants (e.g. 6.84% bacteria). Therefore, the bbduk.sh script from BBMap was used to filter and trim the short reads before alignment to the long-read assembly.

**Table 1.** Sequencing statistics of raw reads for *A. andreniformis* and *A. florea*. Shows the number of raw reads (or read pairs for Illumina) and the total number of gigabases generated. For Illumina reads, estimated coverage is based on the number of sequence bases and an assumed genome size of ∼230 Mb. For ONT long reads, estimated coverage is based on the mean coverage reported by Flye during the initial assembly step. All reads have been uploaded to NCBI SRA and can be downloaded using the listed accession numbers.

| Species (sample) | Sequencing type | Number of raw reads | Number of bases | Estimated coverage | SRA accession |
| --- | --- | --- | --- | --- | --- |
| *A. andreniformis* (Aa1SG1) | ONT long-read | 11,909,664 | 54.0 Gb | 235x | SRR32174174 |
|  | Illumina short-read | 54,425,614 | 16.4 Gb | 71x | SRR32174718 |
| *A. florea* (Af1SG1) | ONT long-read | 10,129,152 | 42.6 Gb | 176x | SRR32174175 |
|  | Illumina short-read | 50,980,281 | 15.4 Gb | 67x | SRR32174721 |

For *A. florea*, we generated 42.6 Gb of long reads using ONT sequencing (read length N50 of 6,132 bp; mean read quality of 15.7). This amount gave us a coverage estimate of 185x based on the assumed genome size of ∼230 Mb. De novo assembly using Flye (Kolmogorov et al. 2019) resulted in a reported mean coverage of 176x, slightly less than our initial estimate. Whole genome sequencing using Illumina yielded 15.4 Gb (50,980,281 read pairs; mean quality score of 38.4), producing an estimated short read coverage of 67x, which we used for base-level polishing of the long-read assembly. As observed with *A. andreniformis*, the ONT reads for *A. florea* showed <0.01 contaminants, whereas the Illumina reads contained 2.47% bacteria and required trimming prior to alignment. Final assemblies were additionally screened using NCBI’s Foreign Contamination Screen (Astashyn et al. 2024) to remove any remaining vector or microbial contaminants before submission to GenBank.

### Assembly quality and completeness

We generated high-quality genome assemblies for *A. andreniformis* and *A. florea* using ONT long-read sequencing, followed by Illumina short-read polishing. The final *A. andreniformis* assembly (Aa1SG1) consisted of 333 contigs totaling 216.8 Mb, a contig N50 of 5.0 Mb, and a GC content of 33.7% (**Table 2**). The *A. florea* assembly (Af1SG1) had a slightly larger total length of 221.6 Mb, with 601 contigs and a contig N50 of 4.3 Mb. Compared to the previously published *A. florea* assembly, Aflo_1.1 (Fouks et al. 2021), which was sequenced using 454 shotgun sequencing, our assemblies show significant improvements in contiguity and completeness. The Aflo_1.1 assembly contains 6,983 scaffolds and 18,378 contigs, with a contig N50 of 24.9 kb–orders of magnitude lower than our assemblies (**Table 2**).

**Table 2.** Assembly contiguity statistics for the *A. andreniformis* and *A. florea* genomes compared to previously published *Apis* genomes. Assembly statistics for our genomes (Aa1SG1 for *A. andreniformis* and Af1SG1 for *A. florea*) were generated using QUAST on the polished assemblies. For the published assemblies (Aflo_1.1 and Amel_HAv3.1), assembly statistics were obtained from their publicly available NCBI genome pages.

| Assembly | Sequencing technology | Total ungapped length | GC content | No. of contigs | Contig N50 | Contig L50 |
| --- | --- | --- | --- | --- | --- | --- |
| Aa1SG1 | ONT + Illumina for polishing | 216.8 Mb | 33.7% | 333 contigs | 5.0 Mb | 14 |
| Af1SG1 | ONT + Illumina for polishing | 221.6 Mb | 33.7% | 601 contigs | 4.3 Mb | 16 |
| Aflo_1.1 | 454 shotgun sequencing | 211.3 Mb | 33.5% | 6,983 scaffolds & 18,378 contigs | scaffold N50: 2.9 Mb, contig N50: 24.9 kb | scaffold L50: 22, contig L50: 2,398 |
| Amel_HAv3.1 | PacBio; 10X Chromium; Phase Genomics HiC; Bionano | 223.9 Mb | 32.5% | 16 chr, 176 scaffolds & 227 contigs | scaffold N50: 13.6 Mb, contig N50: 5.4 Mb | scaffold L50: 7, contig L50: 13 |

Genome completeness estimates using BUSCO confirm the high quality of our assemblies. When evaluated against the Hymenoptera dataset, *A. andreniformis* (Aa1SG1) and *A. florea* (Af1SG1) achieved completeness scores of 98.5% and 98.6%, respectively, with low levels of fragmentation (0.6–0.7%) or missing genes (0.8%; **Table 3**). BUSCO analysis using the broader Insecta dataset showed even higher completeness estimates of 99.7% for both species (**Table 3**). K-mer analysis using Merqury further supported the completeness and base-level accuracy of our assemblies.

**Table 3.** Genome completeness statistics for the *A. andreniformis* and *A. florea* genomes compared to previously published *Apis* genomes. BUSCO analysis was performed using the OrthoDB v10 datasets for Hymenoptera (5,991 genes) and Insecta (1,367 genes), with results shown below. Additional analyses using datasets for Arthropoda (1,013 genes) and Metazoa (954 genes) are summarized in **Supp Table 1**.

| Assembly | Complete<br>[single, duplicated] | Fragmented | Missing |
| --- | --- | --- | --- |
| **HYMENOPTERA** |  |  |  |
| Aa1SG1 | 98.5% [98.4%,0.1%] | 0.7% | 0.8% |
| Af1SG1 | 98.6% [98.5%,0.1%] | 0.6% | 0.8% |
| Aflo_1.1 | 96.8% [96.6%,0.2%] | 1.8% | 1.4% |
| Amel_HAv3.1 | 98.8% [98.6%,0.2%] | 0.3% | 0.9% |
| **INSECTA** |  |  |  |
| Aa1SG1 | 99.7% [99.7%,0.0%] | 0.1% | 0.2% |
| Af1SG1 | 99.6% [99.6%,0.0%] | 0.1% | 0.3% |
| Aflo_1.1 | NA | NA | NA |
| Amel_HAv3.1 | NA | NA | NA |

Since Illumina short reads were used exclusively for polishing rather than assembly, the k-mer completeness estimates primarily reflect how well the short-read data aligns with the final genomes. K-mer completeness was estimated at ∼93% for both species (**Supp Table 2, Supp Figures 1–2**), suggesting that some genomic regions, such as highly repetitive sequences, may be underrepresented in the short-read data. However, the low estimated per-base error rates (∼0.0002, **Supp Table 2**) indicate a high base-level accuracy, consistent with the strong BUSCO scores.

Finally, repeat analysis identified a diverse set of repetitive elements (**Supp Tables 3 & 4**), with total repeat content estimated at ∼6% for both bee species. This content is consistent with previous findings in *A. dorsata* (Oppenheim et al. 2020) and *A. mellifera* (Wallberg et al. 2019). Altogether, our results indicate that these genome assemblies represent the highest-quality *Micrapis* dwarf honey bee genomes published to date.

### Gene annotation

RNA from two pupae per species was extracted and sequenced using a combination of ONT long-read cDNA sequencing and Illumina short-read RNA-seq. For *A. andreniformis*, we generated 70,960,146 ONT long reads with read length N50s exceeding 1 kb (1,178 bp N50 for sample Aa1SG2; 1,145 bp N50 for sample Aa1SG3; **Supp Table 5**), along with 25.6 million high-quality Illumina read pairs. Similarly, for *A. florea*, we obtained 34,084,378 Nanopore reads with read length N50s of 1,098 bp (sample Af1SG2) and 196 bp (sample Af1SG3, excluded from downstream analyses), as well as 21.1 million Illumina read pairs (**Supp Table 5**).

Previous annotations for *A. florea* (Fouks et al. 2021) and *A. mellifera* (Wallberg et al. 2019) identified 12,573 and 12,398 genes, respectively. Our BRAKER3 annotation yielded similar numbers, with 12,232 putative genes for *A. andreniformis* and 12,597 genes for *A. florea* (**Table 4**). BUSCO analysis of the predicted proteomes identified 97.8% of orthologs using the Hymenoptera database and 99.5% of orthologs using the broader Insecta database (**Table 5**), indicating that our annotation successfully captured nearly all expected genes for these species.

**Table 4.** Summary statistics of BRAKER3 gene annotation for both genome assemblies.

| Feature | Aa1SG1 | Af1SG1 |
| --- | --- | --- |
| Number of genes | 12,232 | 12,597 |
| Number of transcripts | 16,282 | 16,689 |
| Number of exons | 111,058 | 112,627 |
| Number of introns | 94,776 | 95,938 |
| Mean gene length (bp) | 7959 | 7455 |
| Mean exon length (bp) | 253 | 254 |
| Mean intron length (bp) | 1396 | 1336 |

**Table 5.** BUSCO completeness statistics for the predicted proteomes of *A. andreniformis* and *A. florea*, assessed using OrthoDB v10 datasets for Hymenoptera (5,991 genes) and Insecta (1,367 genes). Results indicate the percentage of complete, fragmented, and missing orthologs in each assembly, with complete genes further categorized as single-copy or duplicated.

| Proteome | Complete<br>[single, duplicated] | Fragmented | Missing |
| --- | --- | --- | --- |
| **HYMENOPTERA** |  |  |  |
| Aa1SG1 | 97.8% [71.2%,26.6%] | 0.6% | 1.6% |
| Af1SG1 | 97.8% [71.5%,26.3%] | 0.6% | 1.6% |
| **INSECTA** |  |  |  |
| Aa1SG1 | 99.5% [76.2%,23.3%] | 0.0% | 0.5% |
| Af1SG1 | 99.5% [76.1%,23.4%] | 0.1% | 0.4% |

Notably, we observed a relatively high proportion of duplicated BUSCO genes (∼23–26%; **Table 5**), which may reflect uncollapsed heterozygosity, as these diploid female bee genomes were assembled without Hi-C or haplotype resolution. Some alleles may have been retained as separate sequences rather than phased, leading to apparent gene duplications. However, the low fraction of missing BUSCOs (∼0.4–1.6%; **Table 5**) suggests that the assemblies are highly complete, supporting the idea that duplication inflation is due to heterozygosity rather than misassembly and highlighting the need for haplotype-aware approaches in future diploid assemblies.

## Conclusion

Here, we report new high-quality reference genome assemblies for the black dwarf bee *A. andreniformis*, sampled from Frankel, Singapore, and the red dwarf bee *A. florea*, sampled from Admiralty, Singapore. These genomes will provide a crucial resource for further biological, population, and evolutionary studies of dwarf honey bees. Our assemblies were generated using a hybrid sequencing approach combining Oxford Nanopore Technology long reads (with R10.4.1 flowcells) and Illumina short reads. While we sequenced diploid female individuals to enable us to call heterozygous sites, given our relatively modest contig N50 values (5.0 Mb for *A. andreniformis*; 4.3 Mb for *A. florea*), we did not conduct haplotype resolution for these initial assemblies. Ultra-long reads and/or Hi-C data would be necessary to achieve haplotype-resolved and chromosome-level assemblies. Future work sequencing additional individuals from other populations will be critical for developing a comprehensive catalog of genomic variation, including structural variants.

A key strength of our approach is its efficiency, as it required only a single pupa per species to generate genome assemblies with over 98.5% completeness. This approach contrasts with previous studies, such as Aflo_1.1 (Fouks et al. 2021) and Amel_HAv3.1 (Wallberg et al. 2019), which pooled multiple adult drones to achieve similar assembly quality. Our results underscore the potential of using hybrid sequencing to study rare or endangered species, such as these dwarf honey bees, where access to adult specimens is limited or unavailable and even pupae are considered extremely precious.

## Data availability

All raw sequencing data used for genome and transcriptome sequencing are available on the NCBI Sequence Read Archive (www.ncbi.nlm.nih.gov/sra) under the following BioProjects:

### Apis andreniformis

1. Genome sequencing and assembly; NCBI BioProject PRJNA1217036
2. Transcriptome sequencing; NCBI BioProject PRJNA1217108

### Apis florea

1. Genome sequencing and assembly; NCBI BioProject PRJNA1217041
2. Transcriptome sequencing; NCBI BioProject PRJNA1217110

Genome assemblies have been deposited in GenBank under accession GCA_048593525.1 for *Apis andreniformis* and GCA_048593485.1 for *Apis florea*. Gene annotations generated using BRAKER3 are available on Zenodo (https://doi.org/10.5281/zenodo.15048194). Source code and workflows are available on GitHub (https://github.com/atmaivancevic/Micrapis_genome_project).

## Conflict of interest

The authors declare no competing interests.

## Funding

MS and SDR were funded by grant ID AP23PPQS&T00C157, “Study of Honey Bee Pest Diversity to Support Development of Emergency Response Plan”, from the United States Department of Agriculture Animal Plant Health Inspection Service. AI, HA, OJ, and EBC were funded by NIGMS grant ID 2R35GM128822.

## Notes

https://www.ncbi.nlm.nih.gov/bioproject/?term=PRJNA1217036

https://www.ncbi.nlm.nih.gov/bioproject/?term=PRJNA1217108

https://www.ncbi.nlm.nih.gov/bioproject/?term=PRJNA1217041

https://www.ncbi.nlm.nih.gov/bioproject/?term=PRJNA1217110

https://www.ncbi.nlm.nih.gov/datasets/genome/GCA_048593525.1/

https://www.ncbi.nlm.nih.gov/datasets/genome/GCA_048593485.1/

https://zenodo.org/records/15048194

### Competing Interest Statement

The authors have declared no competing interest.

