## Supplementary Material for "De novo genome assemblies of the dwarf honey bee subgenus *Micrapis*: *Apis andreniformis* and *Apis florea*"

**Supp Table 1.** BUSCO completeness statistics for *A. andreniformis* and *A. florea* genomes, assessed using OrthoDB v10 datasets for Arthropoda (1,013 genes) and Metazoa (954 genes). Results include the percentage of complete, fragmented, and missing orthologs in each assembly.

| **Assembly** | **Complete<br>[single, duplicated]** | **Fragmented** | **Missing** |
|---|---|---|---|
| **ARTHROPODA** | | | |
| Aa1SG1 | 99.5% [99.5%,0.0%] | 0.3% | 0.2% |
| Af1SG1 | 99.7% [99.7%,0.0%] | 0.2% | 0.1% |
| **METAZOA** | | | |
| Aa1SG1 | 99.5% [99.4%,0.1%] | 0.2% | 0.3% |
| Af1SG1 | 99.5% [99.4%,0.1%] | 0.2% | 0.3% |

**Supp Table 2.** K-mer analysis statistics for the *A. andreniformis* (Aa1SG1) and *A. florea* (Af1SG1) genome assemblies, generated using Meryl and Merqury. K-mer completeness (%) represents the percentage of short-read k-mers in the genome assembly. Quality value (QV) is a log-scaled metric that quantifies the base-level accuracy of the assembly, with higher values indicating lower error rates. The estimated per-base error rate is derived from the QV score using the formula 10^(-QV/10). It approximates the average base-level sequencing error in the final assembly.

| **Metric** | **Aa1SG1** | **Af1SG1** |
|---|---|---|
| K-mer completeness (%) | 93.90 | 93.28 |
| Total k-mers in Illumina short reads | 213,912,087 | 214,589,584 |
| Total k-mers in genome assembly | 200,856,332 | 200,167,457 |
| Quality value (QV) | 36.47 | 36.34 |
| Estimated per-base error rate | 0.000225 | 0.000232 |

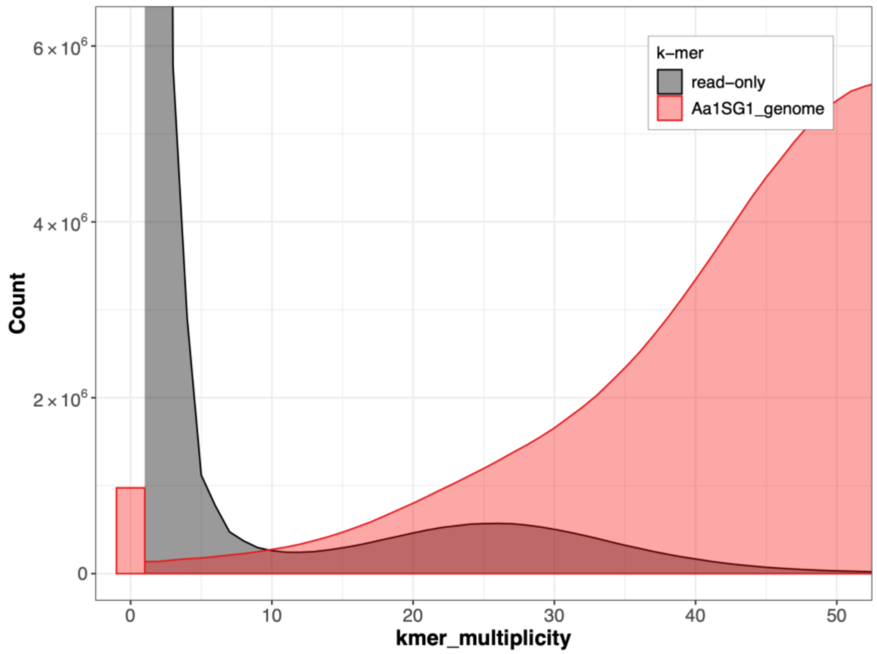


**Supp Figure 1.** K-mer spectra plot for the *A. andreniformis* (Aa1SG1) genome assembly. The k-mer spectra plot visualizes the distribution of k-mer multiplicity in the genome assembly (red) compared to the Illumina short-read dataset (gray). The x-axis represents k-mer multiplicity (how many times a k-mer appears in the dataset), while the y-axis shows the total count of k-mers at each multiplicity level. The "read-only" (gray) peak at multiplicity 1 represents sequencing errors or k-mers not incorporated into the assembly. The shared regions between the short-read dataset and the genome assembly indicate well-represented genomic content.

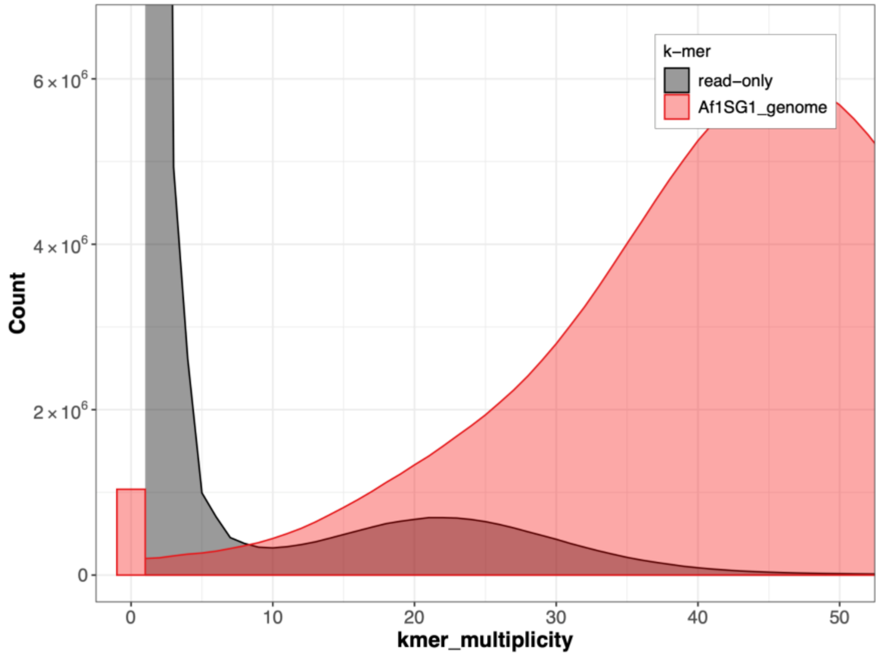


**Supp Figure 2.** K-mer spectra plot for the *A. florea* (Af1SG1) genome assembly.

**Supp Table 3.** RepeatMasker results for *A.andreniformis* (Aa1SG1) genome assembly.

| Repeat type | Number of elements | Length (bp) | Percentage of genome |
|---|---|---|---|
| **Retroelements** | 11960 | 870530 | 0.40 |
| SINEs | 2181 | 132743 | 0.06 |
| Penelope | 1381 | 114731 | 0.05 |
| LINEs | 5568 | 396746 | 0.18 |
| L2/CR1/Rex | 313 | 15294 | 0.01 |
| R1/LOA/Jockey | 266 | 18069 | 0.01 |
| R2/R4/NeSL | 47 | 2418 | 0.00 |
| RTE/Bov-B | 677 | 31984 | 0.01 |
| L1/CIN4 | 1405 | 77509 | 0.04 |
| LTR elements | 4211 | 341041 | 0.16 |
| BEL/Pao | 519 | 101198 | 0.05 |
| Ty1/Copia | 95 | 18356 | 0.01 |
| Gypsy/DIRS1 | 1251 | 81658 | 0.04 |
| Retroviral | 1992 | 108465 | 0.05 |
| **DNA transposons** | 10326 | 763391 | 0.35 |
| hobo-Activator | 2872 | 180969 | 0.08 |
| Tc1-IS630-Pogo | 1310 | 175046 | 0.08 |
| MULE-MuDR | 481 | 25487 | 0.01 |
| PiggyBac | 677 | 103825 | 0.05 |
| Tourist/Harbinger | 595 | 29740 | 0.01 |
| **Rolling-circles** | 598 | 52915 | 0.02 |
| **Unclassified** | 3672 | 257739 | 0.12 |
| **Total interspersed repeats** | | 2006391 | 0.93 |
| **Small RNA** | | 134565 | 0.06 |
| **Satellites** | | 33319 | 0.02 |
| **Simple repeats** | | 9546812 | 4.40 |
| **Low complexity** | | 2270838 | 1.05 |
| **Total repeat content** | | **13949397** | **6.44** |

**Supp Table 4.** RepeatMasker results for *A. florea* (Af1SG1) genome assembly.

| Repeat type | Number of elements | Length (bp) | Percentage of genome |
|---|---|---|---|
| **Retroelements** | 11494 | 741593 | 0.33 |
| SINEs | 2076 | 129602 | 0.06 |
| Penelope | 1419 | 115215 | 0.05 |
| LINEs | 5395 | 309502 | 0.14 |
| L2/CR1/Rex | 309 | 18074 | 0.01 |
| R1/LOA/Jockey | 280 | 19983 | 0.01 |
| R2/R4/NeSL | 49 | 2230 | 0.00 |
| RTE/Bov-B | 641 | 30838 | 0.01 |
| L1/CIN4 | 1327 | 76619 | 0.03 |
| LTR elements | 4023 | 302489 | 0.14 |
| BEL/Pao | 495 | 74139 | 0.03 |
| Ty1/Copia | 82 | 16660 | 0.01 |
| Gypsy/DIRS1 | 1194 | 80488 | 0.04 |
| Retroviral | 1908 | 104922 | 0.05 |
| **DNA transposons** | 9976 | 674897 | 0.30 |
| hobo-Activator | 2813 | 179961 | 0.08 |
| Tc1-IS630-Pogo | 1248 | 155664 | 0.07 |
| MULE-MuDR | 429 | 21252 | 0.01 |
| PiggyBac | 553 | 72229 | 0.03 |
| Tourist/Harbinger | 533 | 26208 | 0.01 |
| **Rolling-circles** | 606 | 46109 | 0.02 |
| **Unclassified** | 3635 | 265600 | 0.12 |
| **Total interspersed repeats** | | 1797305 | 0.81 |
| **Small RNA** | | 122103 | 0.06 |
| **Satellites** | | 34101 | 0.02 |
| **Simple repeats** | | 9509497 | 4.29 |
| **Low complexity** | | 2257595 | 1.02 |
| **Total repeat content** | | **13676061** | **6.17** |

**Supp Table 5.** Sequencing statistics of ONT cDNA and Illumina RNA-seq reads for *A. andreniformis* and *A. florea*. The number of raw reads (or read pairs for Illumina), total bases generated, and read length N50 are shown. All RNA reads have been uploaded to NCBI SRA and can be downloaded using the listed accession numbers.

| **Species (sample)** | **Sequencing type** | **Number of raw reads** | **Number of bases** | **Read length N50** | **SRA accession** |
|---|---|---|---|---|---|
| *A. andreniformis* (Aa1SG2) | ONT long-read | 31,459,925 | 33.0 Gb | 1,178 bp | SRR32174934 |
| | Illumina short-read | 12,019,283 | 3.6 Gb | 150 bp | SRR32174807 |
| *A. andreniformis* (Aa1SG3) | ONT long-read | 39,500,221 | 39.7 Gb | 1,145 bp | SRR32174933 |
| | Illumina short-read | 13,613,083 | 4.1 Gb | 150 bp | SRR32174806 |
| *A. florea* (Af1SG2) | ONT long-read | 32,850,189 | 31.8 Gb | 1,098 bp | SRR32174867 |
| | Illumina short-read | 10,380,023 | 3.1 Gb | 150 bp | SRR32174810 |
| *A. florea* (Af1SG3) | ONT long-read | 1,234,219 | 252.9 Mb | 196 bp | SRR32174866 |
| | Illumina short-read | 10,732,613 | 3.2 Gb | 150 bp | SRR32174809 |
